# Universal Amplicon Sequencing of North Imperial Valley Wetlands Microbiomes

**DOI:** 10.1101/2022.09.29.509762

**Authors:** Scott Becker, Elaina Graham, Lindsay Sager, Roberto Spreafico, Jay McCarren

**Affiliations:** Viridos; Synthetic Genomics

## Abstract

DNA sequencing of complex microbial communities allows for the classification and quantification of thousands of distinct organisms in diverse environmental niches. We present a three domain “Universal Amplicon” (UA) method to simultaneously amplify DNA from the ribosomal small subunit locus from bacteria, archaea, and eukaryotes (and their organelles) using a single pair of amplification primers. We demonstrate the amenability of the UA to multiplexed Illumina library preparation and MiSeq-based sequencing. We validate the UA by sequencing a commercially available microbial community of known quantitative composition and through direct comparison to a shotgun metagenomics dataset. Following validation, we apply the UA to a time-course study of the wetlands of the Northern Imperial Valley in California and show substantial and variable microbial life in the Salton Sea and nearby waters. We demonstrate that the microbial ecology of the Salton Sea varies on at least a monthly basis and is distinct from the surrounding area. Finally, we contribute an open-source Shiny app for real-time analysis of complex metagenomic communities, with application to this study and far beyond.

## INTRODUCTION

Environmental metagenomic methods seek to characterize communities of living organisms that coexist in diverse locales. These techniques can be broadly classified into two classes: shotgun methods that sequence total microbial community DNA[1–4], and amplicon-based methods in which specific sequences (such as 16S ribosomal RNA or ITS) are amplified via polymerase chain reaction (PCR) and subsequently sequenced[5–8]. Through direct sequencing of the genomic content of an entire community, shotgun metagenomics avoids potential biases introduced by amplification steps, can better identify rare taxa[9–12], and provide information on the functional repertoire of the metagenome[13–15]. Despite the continuing decline in sequencing costs, shotgun metagenomic sequencing remains significantly more expensive than amplicon-based approaches[16]. Generally speaking, amplicon methods are able to reconstruct and approximate relative quantitation of those organisms which are present with far lower DNA sequencing requirements than shotgun methods. These amplicon-based approaches have been shown to reproduce the patterns of alpha and beta-diversity which are observed through shotgun sequencing[12]. However, amplicon methods can only detect organisms containing conserved sequences that are recognized by whatever PCR primers are chosen for the experiment; while there are several good choices available for bacterial 16S, eukaryotic 18S, and the Internal Transcribed Spacer (ITS), the development of improved primers is still an area of active research [17–19].

For complex microbial communities that are expected to contain significant representation from multiple domains of life (e.g., diverse bacteria, phototrophic eukaryotes, microbial grazers, etc.), most existing published primers are insufficient to amplify DNA from all community members [20, 21]. While some previous studies have separately amplified sequences for prokaryotic and eukaryotic organisms using multiple sets of primers[22–24], multi-amplicon approaches have a limited ability to quantitatively compare taxa abundances across different primers. There are relatively few published reports of “universal” primers that amplify a more comprehensive subset of living organisms. Amaral-Zettler and colleagues report on the use of “universal” primers[25, 26] but subsequent use of these primer pairs by these authors, or others, appears limited. Two recent publications by Fuhrman et al. show that a single primer pair can meaningfully capture the microbial diversity from all three domains of life in a single amplicon [20, 27].

Previous amplicon and shotgun based metagenomic experiments have shown a fascinating diversity of living organisms in wildly different environmental niches[28–31]. Much of this diversity consists of organisms that are refractory to standard cell culture methods, necessitating metagenomics and amplicon-based approaches for study [32]. One relatively exotic and understudied environment is the Salton Sea, a hypersaline, endorheic lake in Southern California. This “sea” is rapidly receding as reduced riverine and agricultural inputs and generally low precipitation are less than the evaporation rate. As the lake recedes, lake-bed sediments (and contaminants contained within [33]) are exposed and susceptible to movement and transport through the air by wind, resulting in negative health consequences for the people who live nearby[34]. The habitat of many migratory bird species that visit the Salton Sea during their annual migrations is shrinking in conjunction with the receding waters and perturbing the environment in unpredictable ways [35, 36]. Amplicon based metagenomics of the 16S rRNA have previously been conducted at the Salton Sea but these samples had low sequencing depth and were not comprehensively analyzed in the literature [37]. At Viridos, we have an R&D biofuel production facility near Calipatria, California and have undertaken a comprehensive environmental monitoring program in that area, including extensive amplicon-based metagenomic sampling of multiple sites in the vicinity of the southern part of the Salton Sea.

In this report we detail a novel set of universal primers with an optimized library preparation protocol (UA) for use in amplicon-based microbial community profiling. We validate this approach by sequencing a mock community of known composition as well as through comparisons to shotgun metagenomics data. Taken together, our validation studies demonstrate that the UA approach can be used to generate an Illumina-compatible NGS library composed of both prokaryotic and eukaryotic markers. We then utilize the UA methodology to better characterize the unique microbial ecology of the Salton Sea and nearby wetlands, generating a robust dataset with a higher depth of sequencing and resolution than previously existed.

## MATERIAL AND METHODS

### Sample Collection and Processing

Water samples were collected monthly from six separate locations, which we also refer to as stations (**Supplemental Table 1**). At each of these locations, surface water samples were collected in duplicate 500mL sterile polycarbonate bottles, transported to the laboratory on ice, and maintained at 4°C. Microbial biomass from each water sample was collected by vacuum filtration onto a 0.22μm polyethersulfone membrane filter (Pall Corporation, Port Washington NY). Depending on sample type, 50-300 ml of sample was filtered to obtain sufficient biomass for DNA extraction and subsequent analysis (**Supplemental Table 1**). Filters, with collected biomass, were stored in sterile tubes containing milling material (Qiagen, catalog number xxx) at −20C prior to gDNA extraction. All sample processing was conducted within 30 hours of sample collection. Two sites (Managed Marsh and the Alamo River) contain two coordinate locations within close proximity as initial sampling locations became inaccessible during the sample period.

### DNA extraction

Filters stored at −20C in bead tubes were brought to room temperature before performing DNA extractions using the DNeasy PowerWater Kit per the manufacturers protocol (Qiagen, catalog number 14900-100-NF). The DNA was quantified by Qubit dsDNA HS kit (Invitrogen, catalog number Q32851) and normalized to a concentration of 7.0 ng/ul in nuclease free water.

### UA PCR

Normalized gDNA samples were arrayed in 96 well plates and amplified by PCR using barcoded primers according to methods originally described by Caporaso et al. [7], which we have modified to incorporate more universally conserved primer binding sites. Wang et al. [38], identified numerous putative primer binding sites that are highly conserved across eukaryota, bacteria, and archaea. For our work we selected two of the predicted “universal” primers with calculated coverage rates generally greater than 90% across all three domains when compared to the SILVA full-length, non-redundant database [39]. The selected primers amplify a product of approximately 335 bp, not including the primers; exact lengths are 328, 345, and 336 bp for *E. coli, S. cerevisiae*, and *H. volcani* respectively. An amplicon of this size is amenable to generating overlapping paired end sequences using Illumina 2×250bp sequencing technology. The conserved sequences used in the primers are Uni_1046_F (5’ – WGGTGBTGCATGGYYGT – 3’) and Uni_1390_R (5’ – GACGGGCGGTGTGTACAA – 3’). PCR began with denaturation at 95C for 10 min, followed by 30 cycles at 95C for 30 s, 54C for 30 s and 72C for 40 s, and concluded with a final extension for 8 min at 72C.

### Library Preparation and Sequencing

PCR amplicons were pooled at an equal volume of 10ul, purified (Macherey Nagel, Nucleospin Gel and PCR Clean-Up Kit), and eluted in a final volume of 30ul. The pooled amplicon library was quantified by Qubit dsDNA high sensitivity kit (Invitrogen) and amplicon size was determined by Agilent Tapestation 2200 (High Sensitivity D5000 ScreenTape Assay). Library denaturation and dilution was performed according to the standard protocol from Illumina (MiSeq System Denature and Dilute Libraries). MiSeq loading conditions were as follows: 4.0pM library and 0.9375pM PhiX library. Custom read and index primers **(Supplemental Table 2)** were added to the Illumina MiSeq cartridge (MiSeq Reagent Kit v2 (500-cycles)) in well 12 (“Uni_Read I”), well 13 (“Uni_Index”) and well 14 (“Uni_Read 2”). All libraries were sequenced 2×250bp on the Illumina MiSeq following the manufacturer’s instructions and multiplexed with between 100 and 200 samples per run which is sufficient to provide greater than 10000 reads per sample.

#### Bioinformatics Analysis

After sequencing the demultiplexed FASTQ files were collected for analysis. FASTQC (v0.11.7)[40] was run to verify data quality. The R package dada2 (v1.8)[41] was used to denoise the reads, assign counts to amplicon sequence variants (ASVs), and assign taxonomy to ASVs. The analysis pipeline follows the structure of the official pipeline tutorial modified with the following settings: learnErrors (nbases = 4e+08, MAX_CONSIST = 40, multithread = T, randomize = F), dada (separately for forward and reverse reads, pool=”pseudo”, multithread=T), mergePairs (minOverlap= 18, maxMismatch=0, justConcatenate=F, trimOverhang=T), removeBimeraDenovo(method=”consensus”), assignTaxonomy(tryRC=T, minBoot=50, taxLevels = c(“Kingdom”, “Phylum”, “Class”, “Order”, “Family”, “Genus”, “Species”)). Following the mergePairs step all sequences outside of the expected range of 200 to 470 bp were dropped as these are indicative of mispriming. ASVs were retained if they contained at minimum 10 reads in at least one sample; less abundant ASVs were excluded from further analysis. ASV counts were then normalized to 10000 reads. The database used for assignTaxonomy was compiled by pulling all available sequences from Phytoref[42], PR2[43], Silva-Mitochondria [39], and the RDPtraining set 16 [44]. We removed duplicates from the merged database, retained up to seven taxonomic assignments for each sequence, and assigned them to the kingdom, phylum, class, order, family, genus, and species levels. Due to variable taxonomic classification schemas across the databases, we manually curated the taxonomy down to (preferentially) the most accurate genus level annotation or alternatively the lowest level at which the sequence could be accurately discriminated. Prior to the benchmarking study against known mock communities, we filtered the results further to remove any ASV that didn’t occur with at least 100 counts in at least 3 samples, allowing us to focus on how well the measured ASV counts track with the known abundance of each species in this standard.

This data was merged into an Phyloseq object (version 1.26.1)[45] for further analysis. The Phyloseq function tax_glom was used for agglomeration. This function relies on the annotated phylogeny of each ASV to group them together at a given level (kingdom, phylum, etc.). ASVs without an annotation at the desired level are dropped. The Phyloseq functions plot_ordination, plot_richness, plot_bars, and plot_heatmap paired with R’s ggplot functions were used to visualize data.

#### Shotgun metagenomics

To compare the phylogenetic coverage of our UA primers we conducted shotgun sequencing in addition to the UA sequencing on a select set of samples (**Supplemental Table 3**), which were made into a 2×150 library and sequenced on a single lane of the illumina HiSeq platform by Genewiz as part of their Whole Genome Metagenomics service in April of 2020. The reads were trimmed with Trimmomatic (version 0.36; parameters: LEADING:3 TRAILING:3 SLIDINGWINDOW:4:15 MINLEN:26)[46] using standard paired-end Illumina adaptors. MetaWrap (v1.2) [47]was used to QC, assemble, and bin each metagenome using default parameters. Due to low shotgun metagenome coverage a high percentage of bins were <50% complete, so we only include read-based classification via Kraken2 (git hash 18f067d; parameters: −l 150), using a database built on 7-21-2020 with the kracken2-build command (--download-library nt) from the NCBI NT database. The fraction of reads assigned to each phylum were compared to the ratios assigned via the UA method. Phyla representing at least 1% of the reads for any sample were retained for analysis. We then refined the quantifications with Bracken (v<u>2.6.0</u>)[48] and used R (ggplot2)[49] for data visualization.

## RESULTS

### The Universal Amplicon (UA) accurately quantifies a mock community DNA standard

We amplified and sequenced a microbial mock community DNA standard that contains both bacteria and eukaryotes (Zymo Research, Irvine CA) 25 times over the course of this analysis. **Figure 1A** demonstrates the ability of the UA amplicon method to reproduce the relative abundance of the microbial community composition for a standard Zymo mock community. The UA primers capture microbial community composition of both bacteria and eukaryotes with good reproducibility and accuracy, but one bias stands out: the abundance of Salmonella is overestimated while the abundance of Listeria is underestimated. Saccharomyces also shows some minor issues with regard to precision having relative abundances more dispersed across runs than other community members but the average across runs indicates it correlates with the expected abundance (**Figure 1B**). This data demonstrates that the UA primers are capable of detecting both prokaryotic and eukaryotic organisms with relative accuracy despite some dispersion in the data across runs.

**Figure 1.**
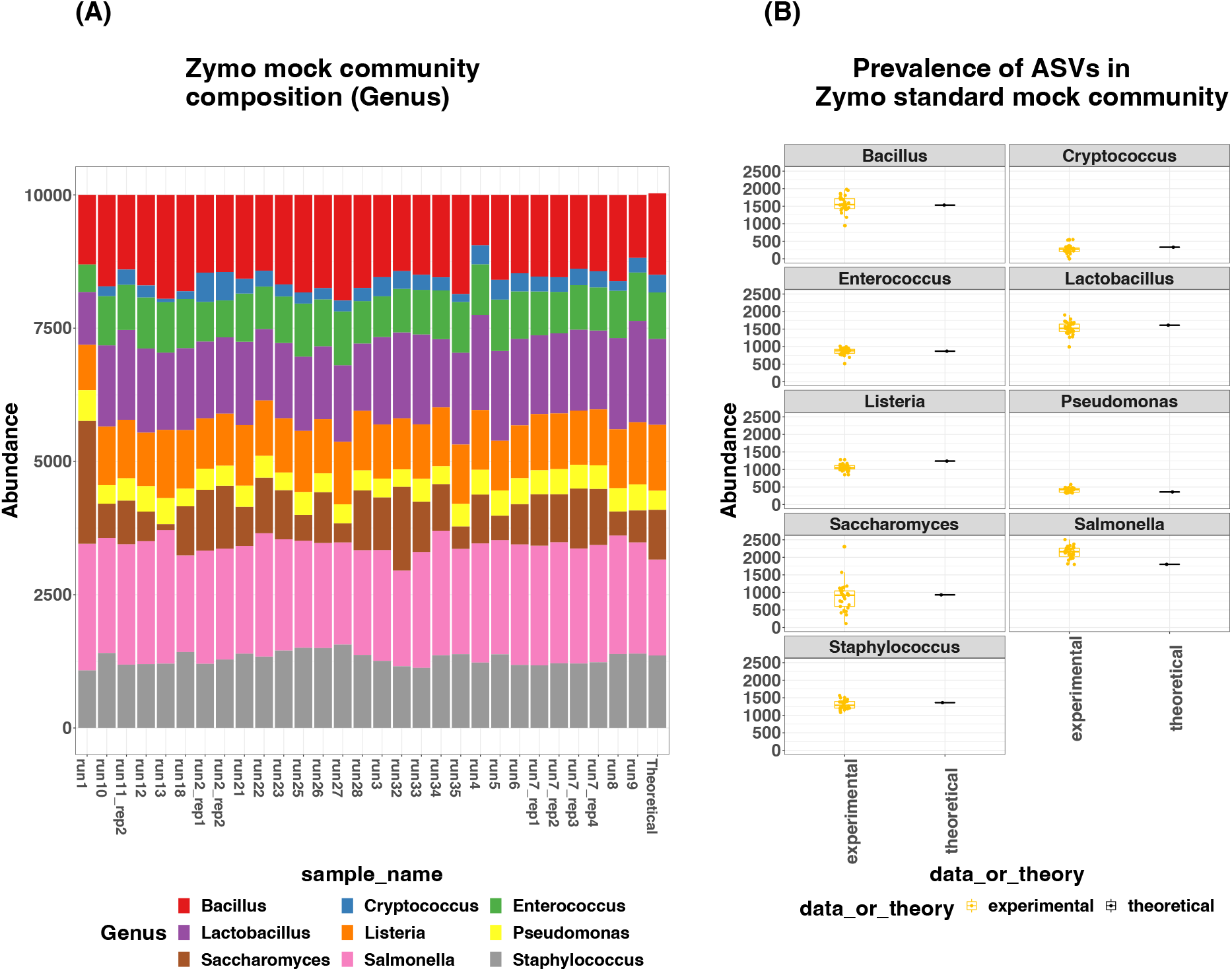
A) Comparison of known composition for the Zymo mock community (far-right bar) compared to 28 sequencing runs using the UA primers described in this paper. Perfect quantitation with the UA method would be an exact match of a given bar to the far-right bar. B) Box and whisker plots showing abundance of each organism within the Zymo mock community over all 28 runs compared to the theoretical or expected abundance of each organism within the mock community.

### UA amplicons show qualitative agreement with shotgun metagenomes at the phylum-level

We performed shotgun metagenomic sequencing of four of our samples (Morton Bay samples from 2018-04-30, 2018-05-30, 2018-06-26, and 2018-07-25) to uncover any potential biases with the UA in real-world data. **Figure 2** shows that most phyla are detected using both the UA and shotgun methods with a high fidelity when comparing the fraction of total reads. Note that the custom UA database (derived from ribosomal marker sequences and explained in the methods) we utilized differentiates between nuclear and chloroplast markers in some cases whereas the NT database used to taxonomically assign reads in the shotgun analysis does not. Some level of discrepancy between the relative abundance of certain phyla in the UA dataset and the shotgun dataset may be due to poor characterization of reads originating from organelles for the shotgun analysis. In addition, marker gene copy numbers (ribosomal and otherwise) per cell vary across lineages [50–52] posing significant quantification challenges when directly comparing relative abundances via shotgun metagenomics and rRNA amplicon relative abundances. Despite these limitations the fraction of total reads from each phylum derived from either shotgun metagenomics or the UA amplicon method overlaps with high fidelity, indicating that the UA primers can generate amplicons across domains without introducing excessive amplification bias.

**Figure 2.**
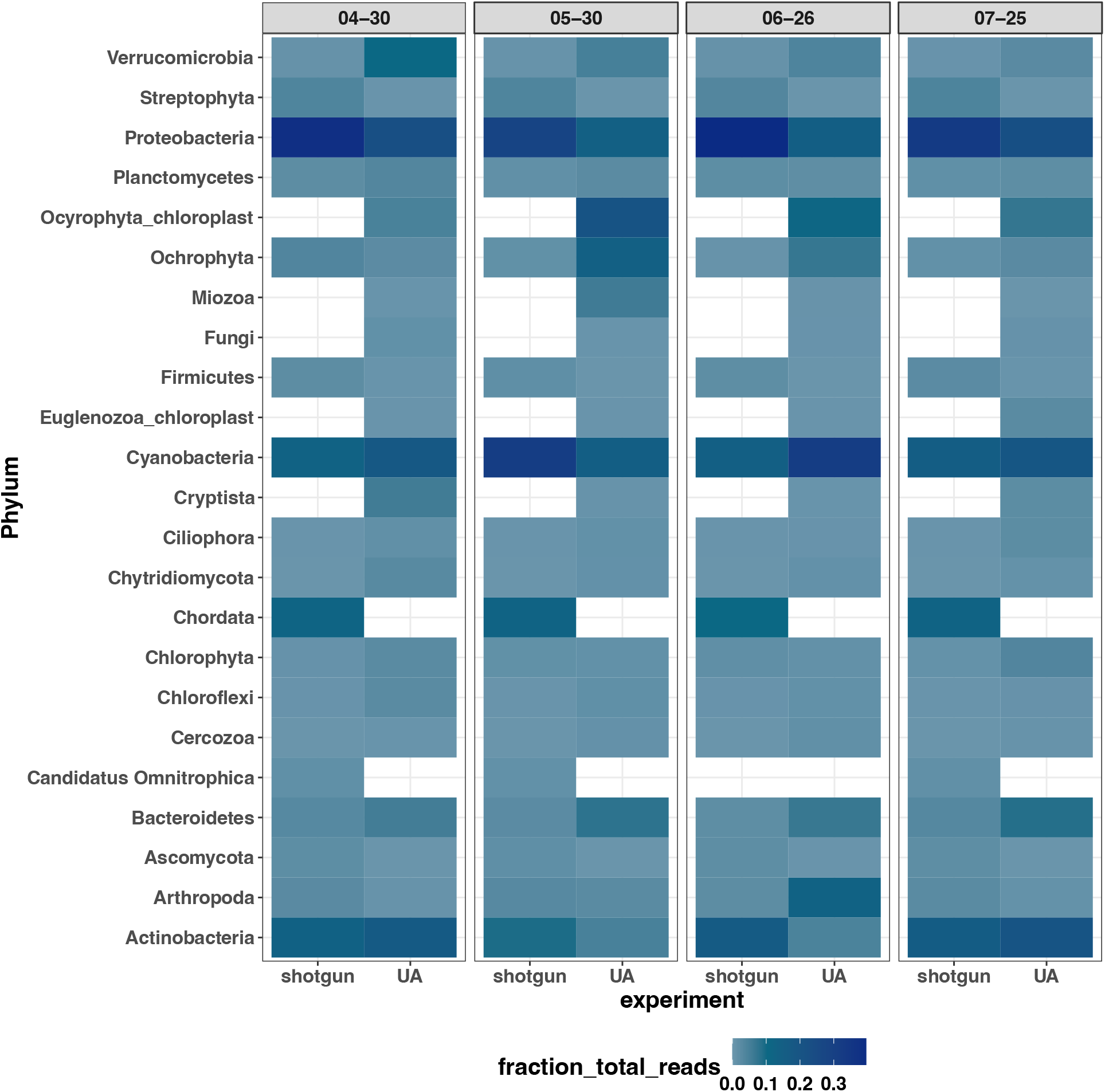
Comparison of shotgun sequencing versus UA characterization of microbial community composition. Heatmap representation of the fraction of sequencing reads corresponding to the given Phyla. Our UA reads contained a significant quantity of reads that could not be classified any more specifically than the Kingdom of Fungi, and hence this kingdom appears adjacent to other phyla.

### California’s Salton Sea and Imperial Valley Wetlands reveal unique communities of diverse organisms

As part of an ongoing environmental monitoring effort associated with farming algae for biofuel production R&D, we sequenced and analyzed a total of 139 water samples (with an average of 39,766 reads per sample) collected approximately monthly from six distinct locations in the North Imperial Valley of California (Error! Reference source not found.) from early 2018 until late 2019. The Salton Sea, which is quickly receding, has become increasingly saline. To date there have been only a handful of analyses of the microbial community within the Salton Sea, including two T-RFLP based studies[53, 54] and two amplicon-based studies [55, 56], of which one only produced data for the prokaryotic community using 16S rRNA primers with low sequencing depth and minimal analysis[56]. The other amplicon study indicates that amplicon sequencing was conducted but meaningful biological data is not presented in the manuscript[55]. Our Salton Sea UA amplicon dataset presented here is the most comprehensive analysis of the microbial community to date indicating that the Salton Sea and nearby environment is home to a variety of distinct organisms with some indication of monthly periodicity in community composition. Non-metric Multidimensional Scaling (NMDS) was used to analyze beta-diversity patterns which demonstrate diversity between samples by measuring similarity vs dissimilarity (**Figure 3A)**. This demonstrated a stark separation of samples collected from the hypersaline Salton Sea and those collected from fresh or brackish waters (e.g Alamo River, N. MacDonald Road, and Managed Marsh/Morton bay), possibly due to different communities of microbes thriving in different salinity conditions.

**Figure 3.**
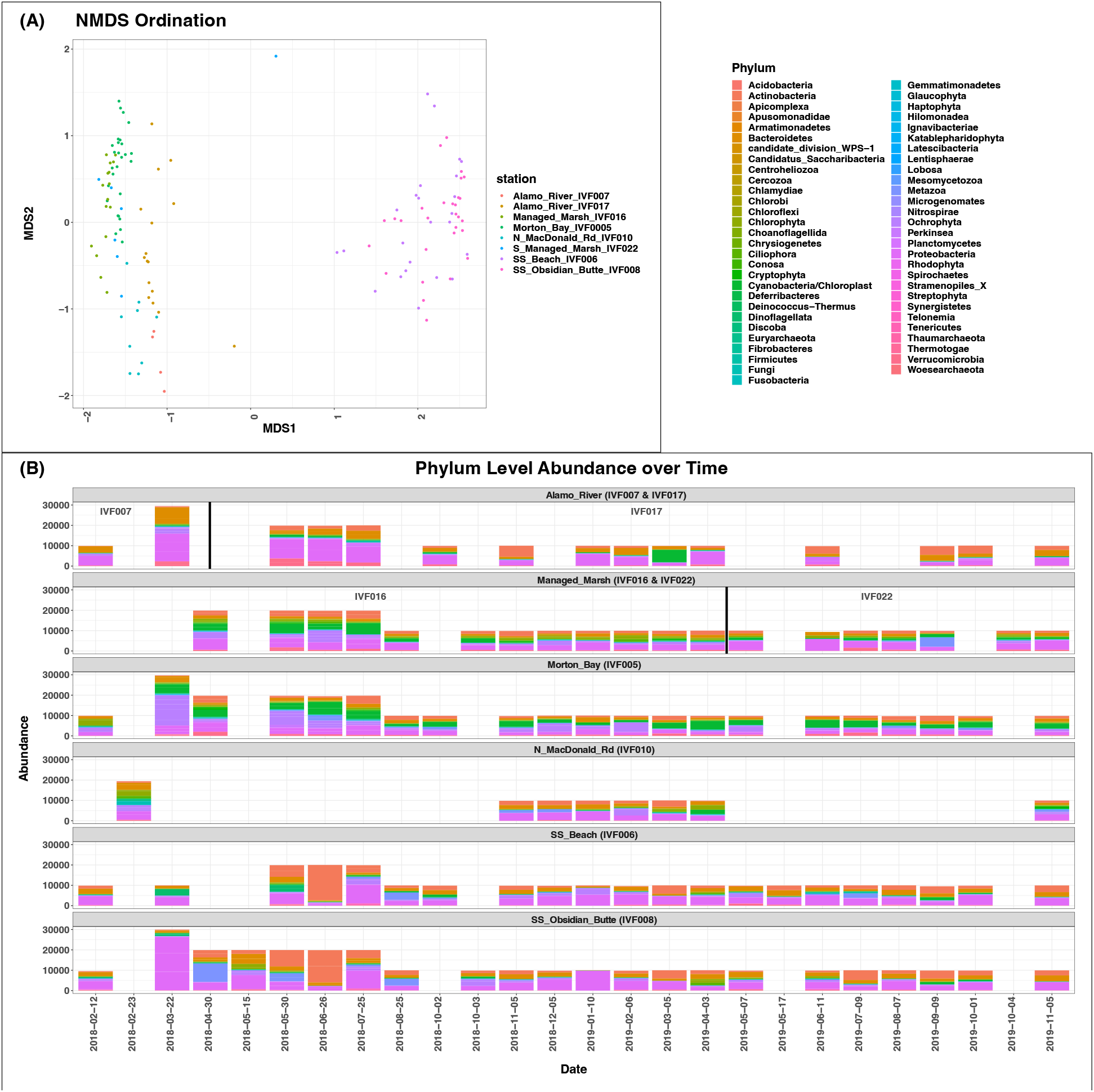
(A) Non-metric multidimensional scaling ordination of each sample. Samples are colored by station ID. (B) Bar plot of ASVs over time at each station agglomerated to the Phylum level. Vertical lines mark where specific sampling locations for the Alamo River and Managed Marsh changed due to previous locations becoming inaccessible.

Within these distinct samples we observe thousands of unique Amplicon Sequence Variants (ASVs). While ASVs are useful in identifying fine scale community level differences, each unique ASV is not necessarily representative of a distinct species because many organisms have multiple rRNA copies (which are assigned to unique ASVs unless they are identical in sequence across the amplicon we used) and individual members of species have slight variations in exact rRNA sequence. Rarefication analysis, which allows us to calculate full sample richness vs observed sample richness indicated asymptomatic behavior in most samples with the exception of early samples from the Alamo River site in 2018 (which was abandoned early in the study due to accessibility issues) **(Supplemental Figure 1)**.

In total we observed over 8000 ASVs and the most prevalent 75 taxa were identified for visualization across samples (**Supplemental Figure 2**). NMDS ordination immediately shows that distinct communities of microorganisms are present in the Salton Sea and the nearby wetlands. The Salton Sea has abundant Trebouxiophyceae, Dinophyceae, and Gammaproteobacteria; whereas the surrounding wetlands have abundant Cyanobacteria. When agglomerated by class these differences are even more pronounced and the non-Salton Sea samples are almost completely devoid of certain classes of organisms (**Figure 3B**).

When we agglomerate to the kingdom level (**Supplemental Figure 3**), we see significant shifts in the prokaryotic/eukaryotic ratio of ASVs across sites and over time, potentially indicating some form of seasonal periodicity, but more sampling over an extended period of time would need to be conducted to confirm this. Phylum level agglomeration shows that in both 2018 and 2019, a dramatic shift from Proteobacteria and toward Actinobacteria was observed in the early summer at the Obsidian Butte sampling location in the Salton Sea. This trend is similarly reflected, somewhat less starkly, at the Salton Sea Beach sampling station. At Obsidian Butte we see that between the extremes of this shift, ASVs assigned to Bacteriodetes, Chlorophyta, Metazoa, and Ochrophyta increase in abundance before being completly overtaken by Actinobacteria. Limited sampling at the Salton Sea Beach during this transition makes it difficult to assess whether this same trend occurs here, but the available sampling suggests this could be a shared trend.

For a deeper analysis, we focus on Obsidian Butte samples from March through June of 2018, retaining all ASVs from these months with more than 10 counts in at least one sample. Alpha diversity indices show a peak in ASV richness in the May 15 sample (**Figure 4A**). At the phylum level of this subset of samples there is a dramatic shift from proteobacteria to actinobacteria (**Figure 4B**). The mid-May sample has a higher abundance of phototrophs (Chlorophyta, Ochrophyta, and dinoflagellates), suggesting a potential spring bloom. The subsequent late May sample shows a significant increase of actinobacteria. A previous study has demonstrated that during a dinoflagellate bloom showed that Gammaproteobacteria, Bacteriodetes, and Alphaproteobacteria were prevalent during the initial bloom stages while Cyanobacteria and Actinobacteria were more prevalent during bloom termination[57]. The April and late May samples also show increases in the Metazoa assigned ASVs (specifically Apocyclops royi, which is a copepod). While extensive grazer activity may be related to drastic shifts in species abundance in microbiomes, we did not conduct size filtering while sampling and the high potential copy number of rRNA amplicons proliferating from a single copepod could produce technical bias, so we cannot make a strong case for grazer-related shifts in this dataset.

**Figure 4.**
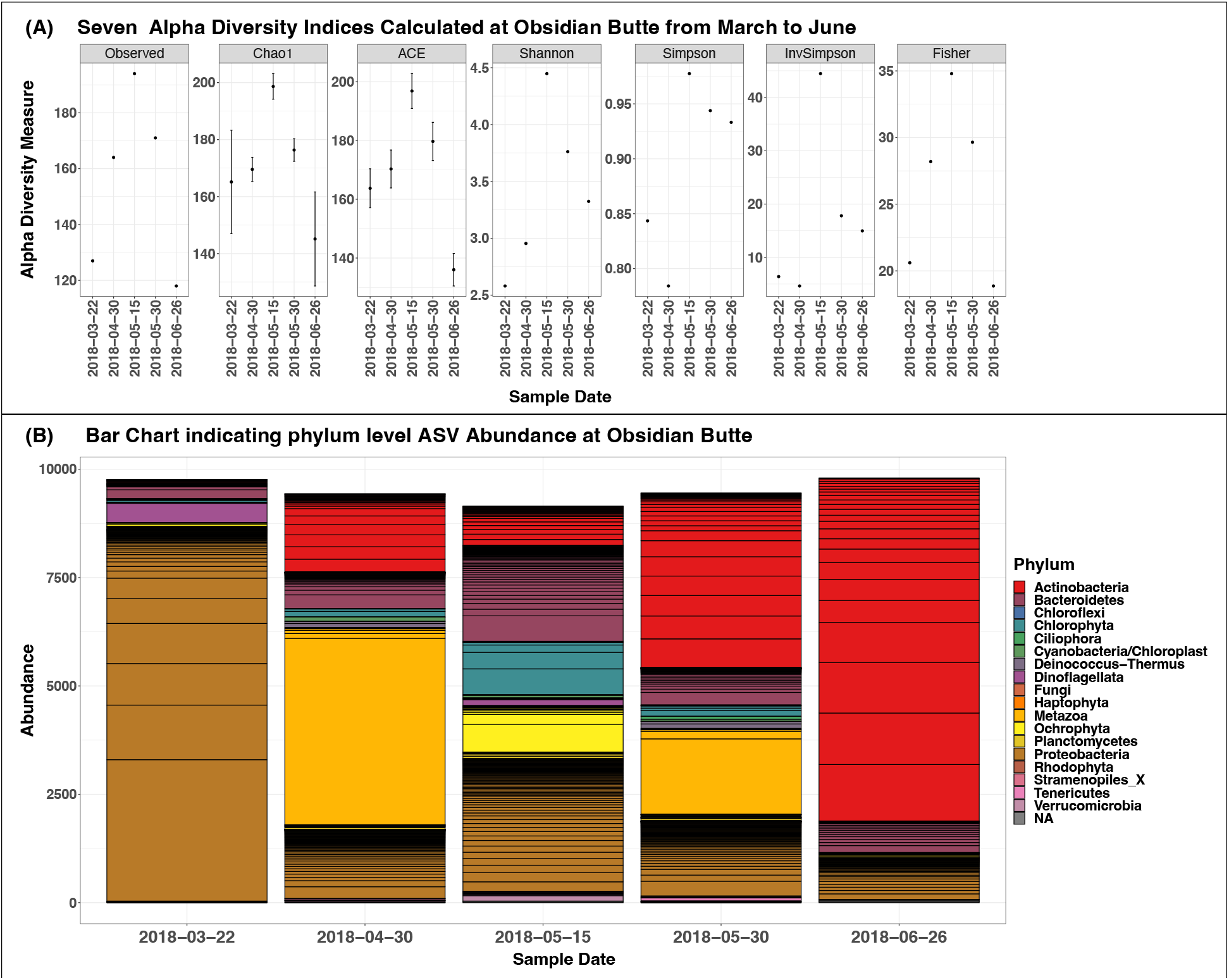
A) Seven Alpha diversity indexes shown side by side to assess evenness and richness of the community. X axis reflects dates of interest and y axis the calculated alpha diversity. The higher the value of y the more diverse the community is. The measures used include Observed diversity index, Chao1 diversity index, ACE diversity index, Shannon diversity index, Simpson diversity index, Inverse Simpson diversity index and Fisher diversity index. (B) ASV abundance at Obsidian Butte from March through June of 2018. ASV abundances were agglomerated to the phylum level.

### Discussion

Through the use of control standards and comparisons to shotgun metagenomics, we have herein demonstrated an efficient, single-amplicon based approach that is able to quantitatively distinguish both prokaryotic and eukaryotic organisms simultaneously through straightforward sequencing on the Illumina MiSeq platform. The data analysis tools for this sort of data are (with the exception of requiring publicly available eukaryotic marker sequences) essentially identical to typical 16s methods. While a very limited number of previous publications have demonstrated some of the same capabilities, we have synthesized a workflow that can be rapidly applied to varied environmental niches for many different purposes.

As a proof of concept, we sequenced the wetlands near the southern portion of the Salton Sea numerous times over many months. While the Salton Sea outwardly appears to be a dying ecosystem, our study of microbial life clearly shows a tremendous and highly variable diversity of microorganisms. While outside the scope of our current study, one could imagine correlating environmental variables with microbial communities to better understand how the ecosystem might change as the Salton Sea continues to recede. Such an effort might also suggest ways in which microbial life could contribute to mitigating the worst effects of the gradual disappearance of the sea.

One limitation of sequencing relatively short (<500 bp) amplicons for phylogenetic characterization of communities is that the level at which many organisms can be uniquely classified tends to be less specific than species. We see this phenomenon in much of the data presented herein, where agglomeration even up to the phylum level is sometimes needed to present clear trends. Looking forward, the Oxford Nanopore platform offers an interesting opportunity to sequence longer amplicons and better distinguish between closely related organisms[58], possibly even at the species level. Due to the form factor, this technology could potentially allow sequencing on site rather than transporting samples significant distances. Prior to implementation across kingdoms of life, new primers for a longer universal amplicon would need to be validated.

Finally, while dedicated researchers in the field, especially the team working on phyloseq, have made great progress in enabling straightforward microbiome data analysis, advanced computational skills are still a barrier to entry into the field. By providing a GUI-based app for microbiome analysis, we hope to allow a wider group of researchers a quicker route into sophisticated, first-class, microbiome data analysis.

## Conclusions

We present a novel method for microbiome amplicon sequencing (UA) that is able to simultaneously amplify DNA from all three kingdoms of life and show that it is in agreement with a known control community and a proven shotgun metagenomics approach. We demonstrate the value and potential of the UA by sequencing water samples in the North Imperial Valley wetlands of California, including the Salton Sea, and show microbial diversity across both space and time in this sensitive geographic area.

## Data and software availability

We have written code for a highly functional R Shiny app for interactive, user-driven analysis of amplicon microbiome data. Inspired by the Shiny-phyloseq app (https://joey711.github.io/shiny-phyloseq/) and the rich capabilities of the phyloseq package, we have coded a real-time user interface for common microbiome analysis and visualization tasks. The app takes a phyloseq object as input and is capable of a variety of analyses, including sample/taxa filtering, tree-building, beta diversity and ordination, bar plots, heatmaps, permanova, and differential abundance. In essence, we wrapped a GUI around what we see as best practices for command-line microbiome analysis in R. The app is a work in progress, and open to contributions from the community at gitlab.com/viridos-asv. A webpage pre-loaded with the data from this study can be accessed at asv.viridos.com. Users should note that some of the analysis results described in this paper require code beyond what is available in this app; we have also posted a R Markdown document that recreates this analysis on gitlab along with the app code.

## Acknowledgments

The authors would like to acknowledge Bhargavi Ramanathan for help gathering samples, Laura de Boer for critical comments on the manuscript, and Rob Brown for inspirational discussions. Environmental sampling activities were performed during work authorized under U.S. EPA TERA Permits R-19-0001 and R-20-0001. We greatly appreciate the assistance and cooperation of our neighbors in Calipatria, CA. Our thanks go to Mr. Walter Slovak, the County of Imperial, the Imperial Irrigation District, CA Department of Fish & Wildlife, BHE Renewables, Controlled Thermal Resources Ltd., and EnergySource Inc. for access to their lands and waters.

## Supplemental Figures

**Supplemental Figure 1:**
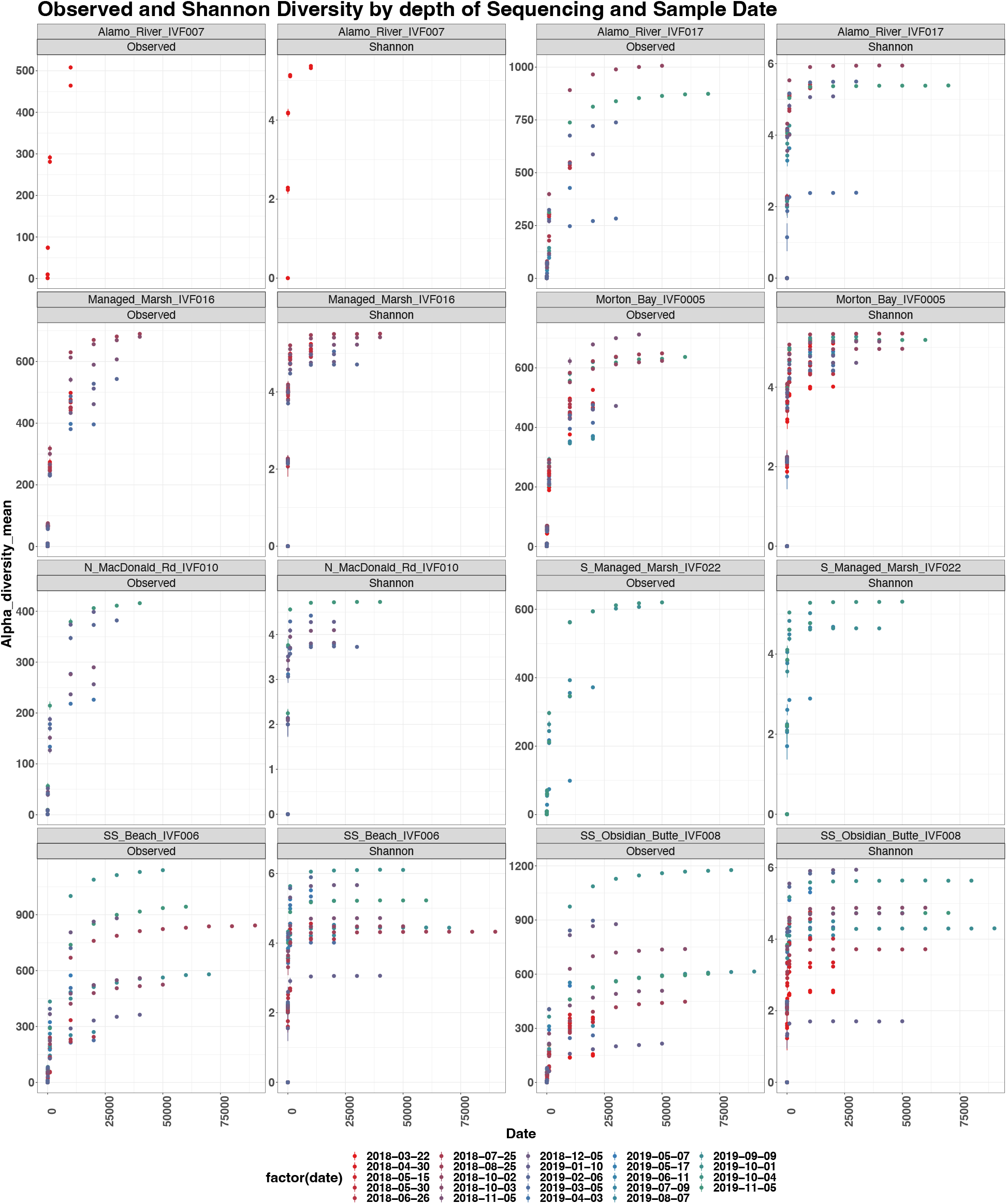
Rarefication analysis across sample sites with predicted, observed, and Shannon diversity calculated across sequencing depth. Points are colored by unique sampling date.

**Supplemental Figure 2:**
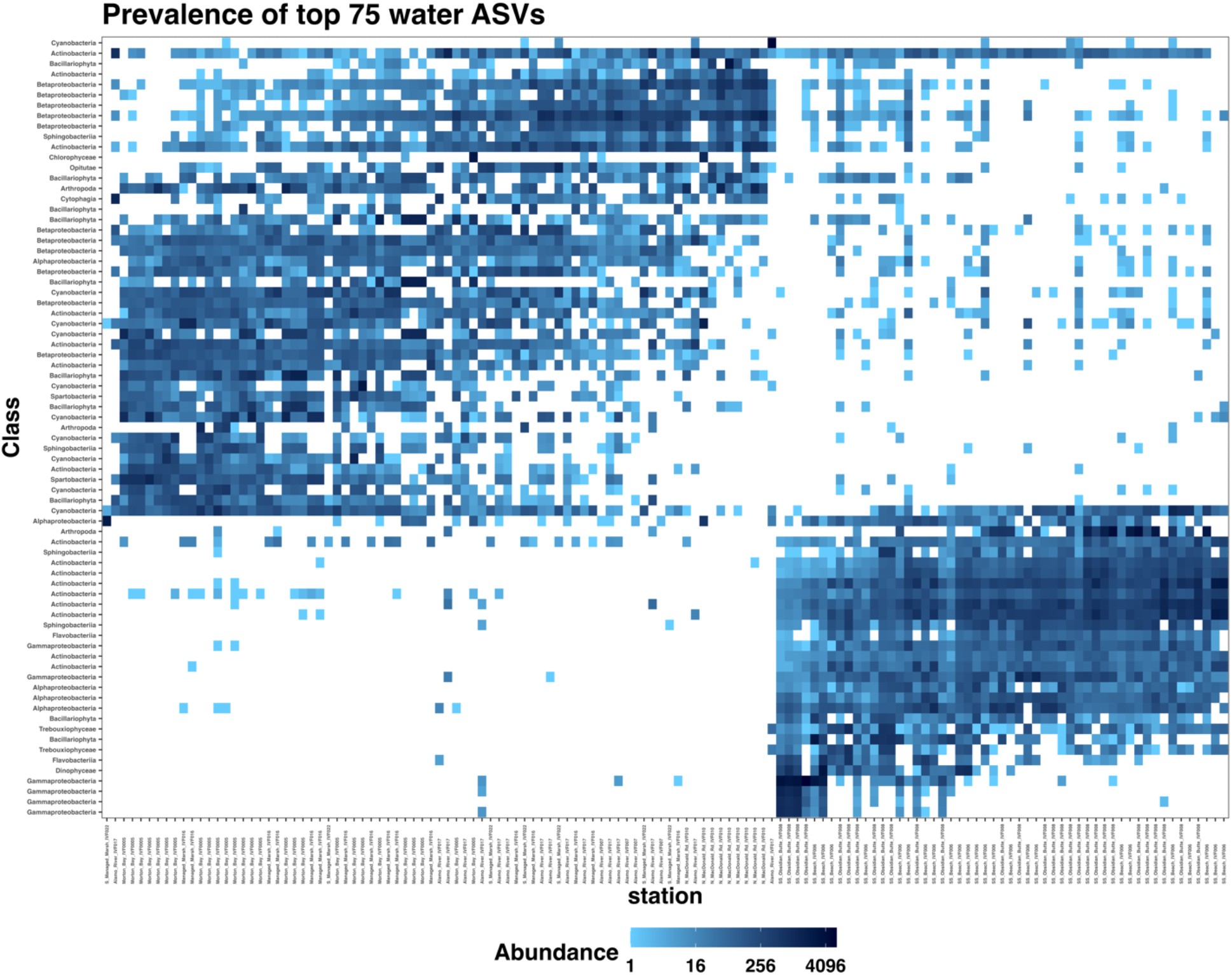
Heatmap of the top 75 ASVs agglomerated to the class level columns sorted by sample name.

**Supplemental Figure 3.**
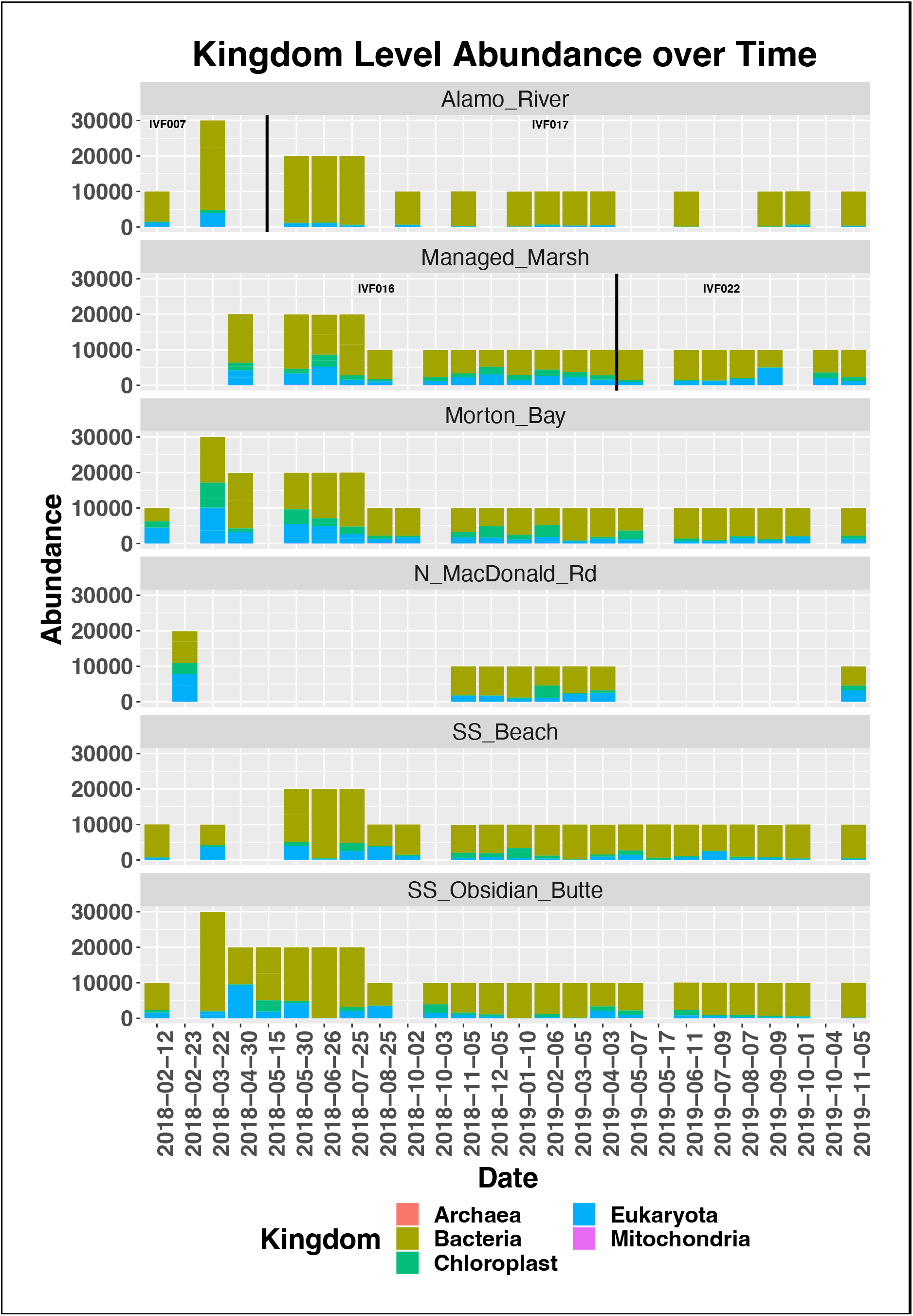
Bar plot of ASVs over time at each station agglomerated to the Kingdom level

**Supplemental Figure 4.**
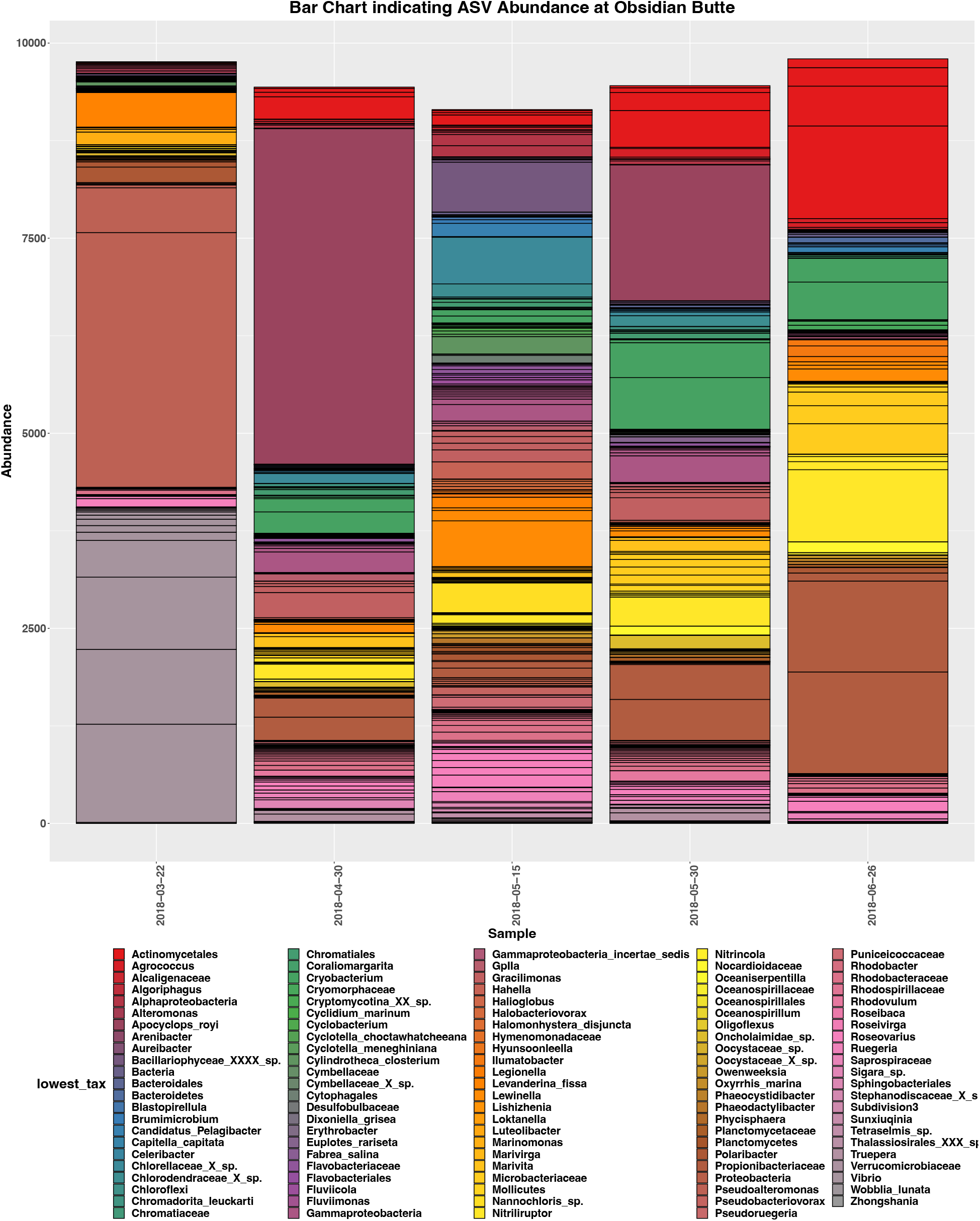
A bar chart showing ASV abundance at Obsidian Butte from March through June. ASV abundances were agglomorated to the lowest level at which a unique classification was possible.

**Supplemental Figure 5.**
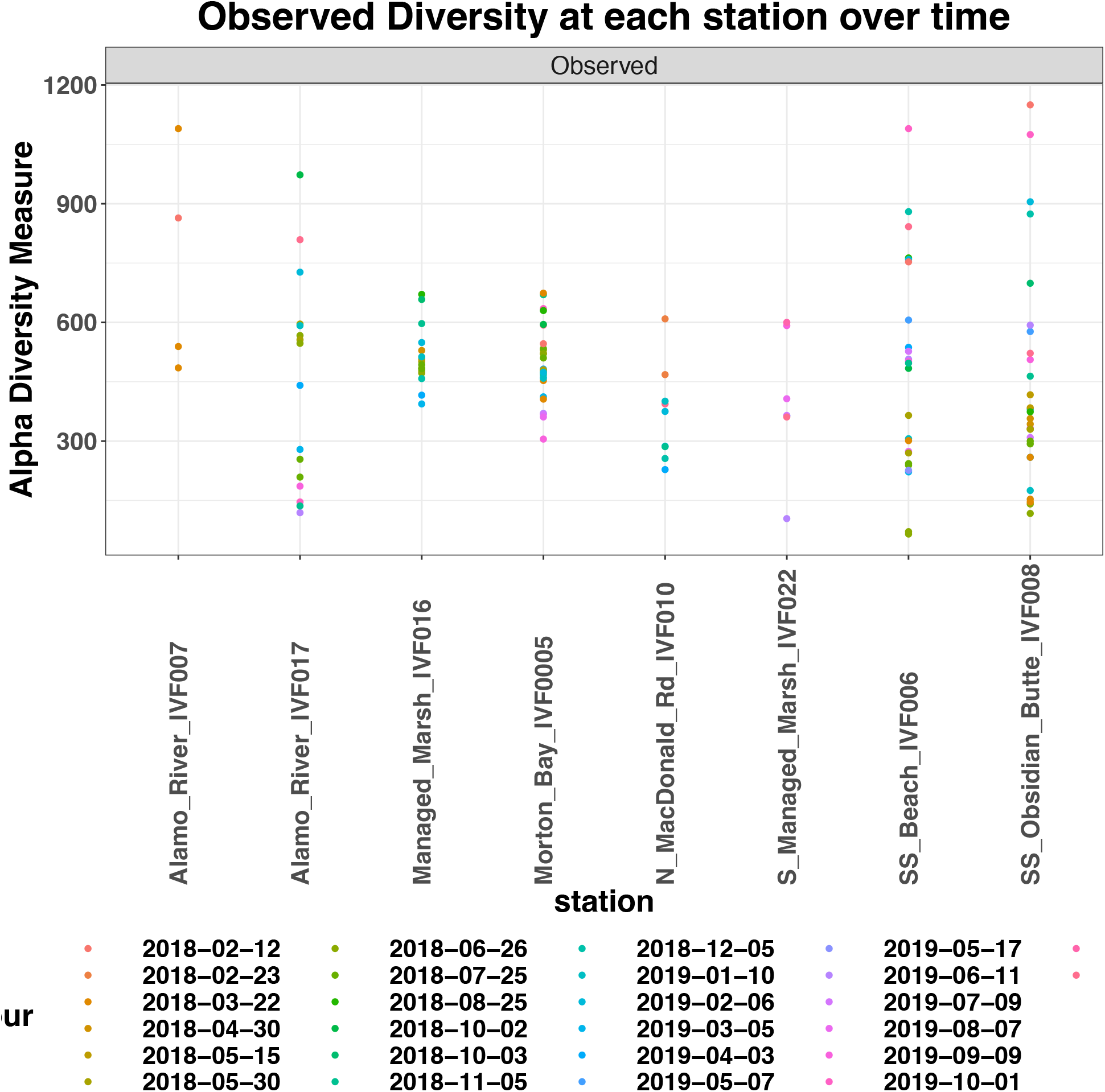
Shannon diversity (calculated using phyloseq’s plot_richness) function across all sample sites.

**Supplemental Figure 6.**
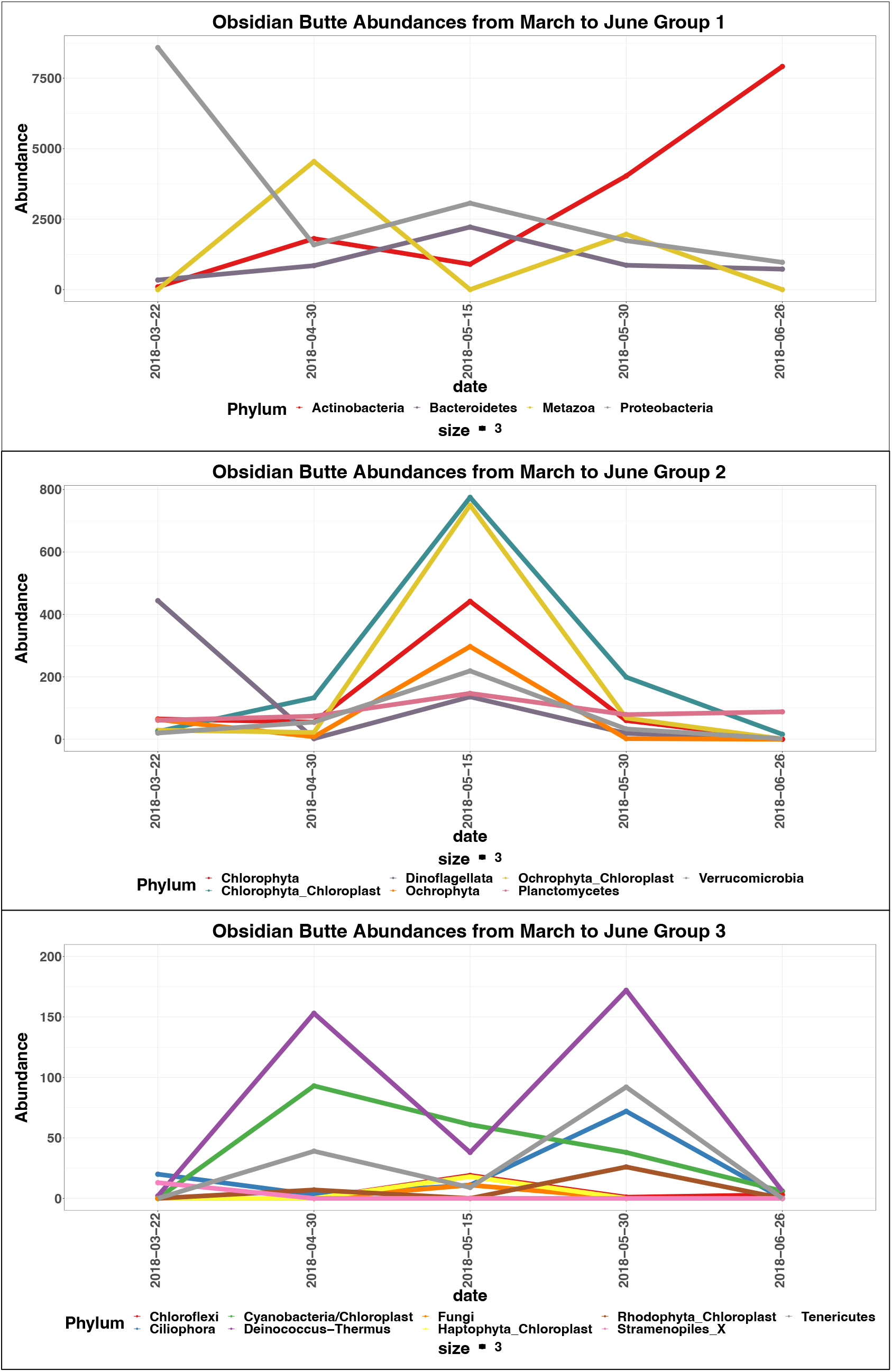
Obsidian Butte Phylum level abundances from March through June of 2018 separated into unique facet groups. Groups were chosen based on known correlations between certain species in the literature and overlapping trends in abundances.

